# Nipsnap1—A Regulatory Factor Required For Long-Term Maintenance of Non-Shivering Thermogenesis

**DOI:** 10.1101/2022.04.29.490099

**Authors:** Yang Liu, Yue Qu, Chloe Cheng, Pei-Yin Tsai, Kaydine Edwards, Siwen Xue, Supriya Pandit, Sakura Eguchi, Navneet Sanghera, Joeva J Barrow

## Abstract

The molecular activation of non-shivering thermogenesis (NST) has strong potential to combat obesity and metabolic disease. However, the mechanisms surrounding the maintenance of NST once it is fully activated, remain unexplored. Here, we present 4-Nitrophenylphosphatase Domain and Non-Neuronal SNAP25-Like 1 (Nipsnap1) as a critical regulator of long-term thermogenic maintenance in brown adipose tissue (BAT). Nipsnap1 localizes to the mitochondrial matrix and increases its transcript and protein levels in response to both chronic cold and β_3_ adrenergic signaling. Through the generation of BAT-specific Nipsnap1 knockout mice (N1-KO), we show that these mice are unable to sustain activated energy expenditure and fail to protect their body temperature in the face of an extended cold challenge. Mechanistically, we demonstrate that Nipsnap1 integrates with lipid metabolism and BAT-specific ablation of Nipsnap1 leads to severe defects in β-oxidation capacity. Our findings identify Nipsnap1 as a potent regulator of long-term NST maintenance.

## INTRODUCTION

The metabolic state of obesity results from the chronic and excessive accumulation of fat either due to overnutrition and/or insufficient energy expenditure (Piché, Tchernof and Després, 2020; Panuganti, Nguyen and Kshirsagar, 2021). It is well-established that obesity is associated with adverse health outcomes including diabetes, cardiovascular diseases, and several types of cancer (Pi-Sunyer, 2009; Chobot *et al*., 2018; Avgerinos *et al*., 2019; Cercato and Fonseca, 2019). Indeed, the global prevalence of obesity in humans has crested to pandemic proportions and it is now classified as the leading cause of preventable death worldwide (Censin *et al*., 2019). Current therapies such as dietary restriction, surgery, or pharmacological interventions are either unsustainable or are limited by adverse side effects (Bal *et al*., 2012; Chang *et al*., 2014; Thom and Lean, 2017; Siebenhofer *et al*., 2021). Therefore, the need for novel therapeutic options is critical. The activation of the brown and beige fat non-shivering thermogenic (NST) program to increase energy expenditure has been conclusively proven to protect against obesity in rodent models and is promising as a molecular treatment option to combat obesity in humans (Rothwell and Stock, 1997;Bartelt *et al*., 2011; Bordicchia *et al*., 2012; K. Y. Chen *et al*., 2013). Indeed, human brown adipose tissue (BAT) can be safely activated by acute cold exposure and is associated with an increase in energy expenditure (Lichtenbelt et al., 2002; Claessens-Van Ooijen et al., 2006; Cypess et al., 2009; van Marken Lichtenbelt et al., 2009; Zingaretti et al., 2009; Celi et al., 2010). While the mechanisms underlying BAT thermogenic activation are increasingly well understood, the mechanisms regarding how NST is sustained, once activated, are poorly characterized and relatively under-explored (Hondares *et al*., 2010; Seale *et al*., 2011; Kleiner *et al*., 2012). This knowledge gap becomes extremely important given the transient nature of BAT activation. As has been shown in human studies, once thermogenic stimuli are removed, BAT is rapidly deactivated and most protective benefits against obesity are lost (Roh *et al*., 2018; Leitner *et al*., 2019). Identifying regulatory factors that are required for long-term thermogenic maintenance will provide additional molecular therapeutic opportunities to maintain the protective benefits of NST in order to combat obesity and associated metabolic diseases. In order to address this problem, we have mapped the BAT mitochondrial proteome in mice under chronic cold challenge conditions to identify unique protein signatures that either become upregulated, or demonstrate sustained upregulation in late-stage thermogenesis—defined as greater than or equal to 7 days of cold exposure. We hypothesize that proteins exhibiting this expression pattern could be involved in the regulation of long-term thermogenic maintenance in BAT.

Here, we have discovered a novel thermogenic factor known as 4-Nitrophenylphosphatase Domain and Non-Neuronal SNAP25-Like 1 (Nipsnap1) as a potent regulator of long-term thermogenic maintenance in BAT. Nipsnap1 is an evolutionarily conserved protein from fly to humans that is expressed predominantly in highly energetic tissues such as the brain and liver (Seroussi *et al*., 1998; Satoh *et al*., 2002; Nautiyal *et al*., 2010; Tummala, Li and Homayouni, 2010; Morgenstern *et al*., 2021). It possesses an N-terminal mitochondrial targeting sequence (MTS) that directs it to the mitochondrial matrix, and previous studies have determined that Nipsnap1 plays a critical role in mitophagy where it functions as a sensor of mitochondrial health and recruits autophagy proteins when mitochondria become damaged (Abudu *et al*., 2019; Fathi, Yarbro and Homayouni, 2021). Other reported functions include its role in pyruvate and branched chain amino acid metabolism (Manisha Nautiyal *et al*., 2008; Ghoshal, Jones and Homayouni, 2014), pain transmission signaling, neurological disorders, carcinogenesis, and the immune response (Okuda-Ashitaka *et al*., 2012; Yamamoto *et al*., 2017). Most of the current research for Nipsnap1 however details these functions in brain or liver tissues but the expression profile and function of Nipsnap1 in thermogenic adipose tissue is unknown. In this study, we provide evidence of Nipsnap1 as a late-stage thermogenic regulatory factor. We show that Nipsnap1 is robustly activated only after chronic cold exposure in BAT. We further demonstrate that loss-of-Nipsnap1 in brown fat depots impairs the sustained activation of the NST program. Mechanistically, ablation of Nipsnap1 in BAT compromises mitochondrial lipid beta-oxidation capacity leading to a significant decline in energy expenditure and an inability to maintain body temperature homeostasis in the face of chronic cold exposure. Thus, we show that Nipsnap1 is a critical regulator of NST through its role in lipid metabolism and may have targeted therapeutic opportunities to ensure the long-term activation of NST.

## RESULTS

### Nipsnap1 has potent late-stage thermogenic properties

To identify proteins thathave the potential to serve as key mediators of late-stage thermogenic maintenance, we isolated mitochondria from interscapular BAT of mice exposed to either a chronic cold (6.5°C) or a thermoneutral (30°C) environment for 8 days and performed TMT-labeled proteomics (Figure 1A and 1B). Of the captured mitoproteome, a total of 85 proteins were strongly induced in chronic cold exposure compared to thermoneutral controls. Candidates were subsequently filtered for proteins with orthologous expression profiles in human BAT leaving only 8 candidates that were completely unannotated for a role in BAT non-shivering thermogenesis (Müller *et al*., 2016). Of these candidates, Nipsnap1 emerged as the most significant based on protein and gene abundance levels (Figure 1C and 1D).

**Figure 1.**
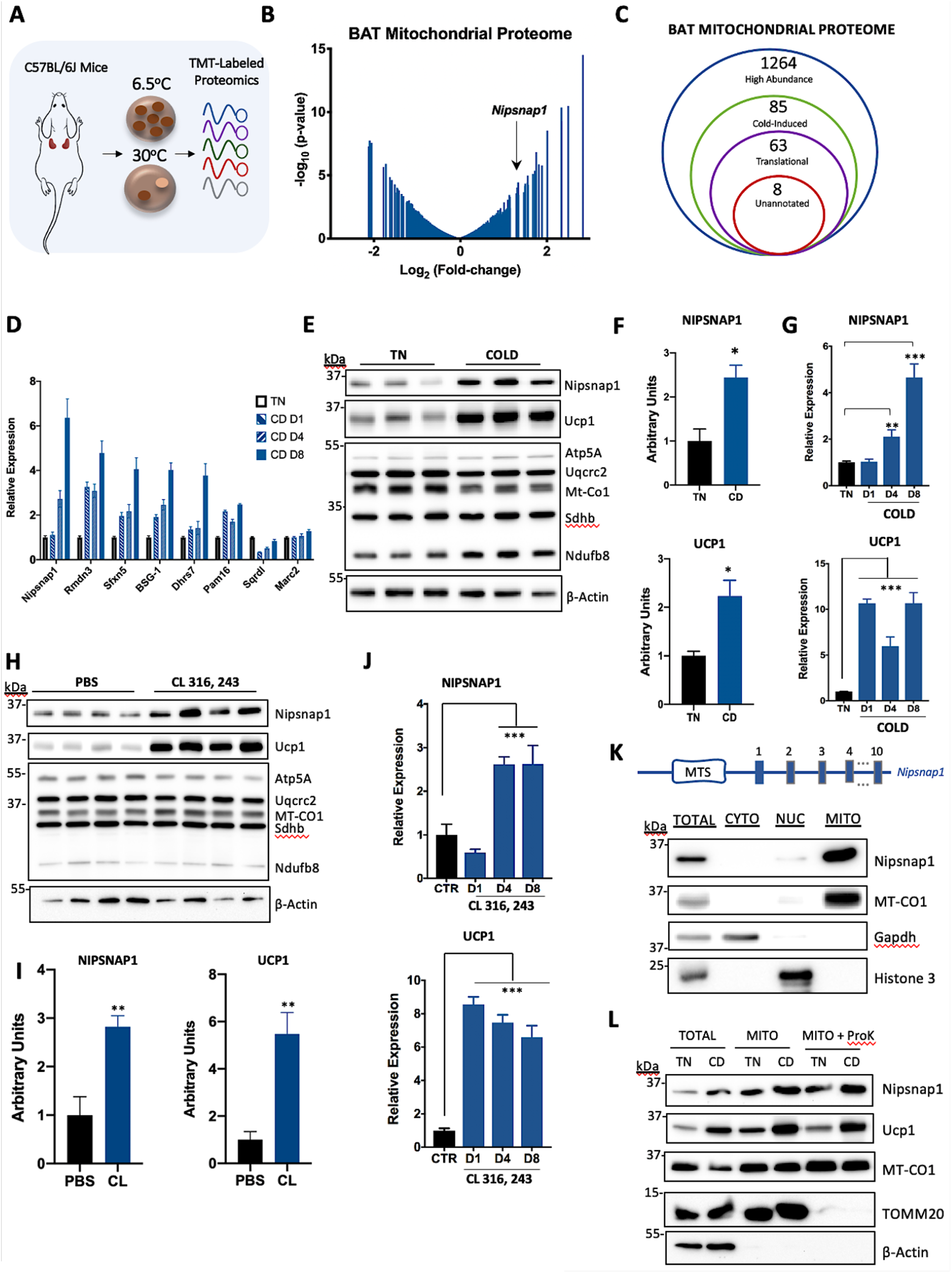
Nipsnap1 displays potent late-stage thermogenic properties. **(A)**Schematic of BAT mitochondrial proteomic analysis in chronic cold compared to TN-exposed wild-type (WT) mice. **(B)** Line volcano plot of the mitochondrial proteins identified in BAT mitochondria (cold vs TN). **(C)** Diagram of filtering process to identify unannotated mitochondrial proteins. **(D)** Relative mRNA expression from WT mice exposed to 1, 4, and 8 days of cold (CD) exposure compared thermoneutral controls (n=3). **(E)** Representative Western blot of BAT tissue from WT mice exposed to TN and Cold for 8 days. **(F)** Densitometry of Nipsnap1 (above) and Ucp1 (below) protein levels normalized to β-Actin from Figure 1E. **(G)** Relative mRNA time-course expression of Nipsnap1 (above) and Ucp1 (below) from WT mice exposed to 1, 4, 8 days of cold exposure compared to a thermoneutral controls (n=5). **(H)** Representative Western blot of BAT tissue from mice treated with CL 316,243 or PBS IP injection for 8 days. **(I)** Densitometry of Nipsnap1 (left) and Ucp1 (right) protein levels normalized to β-Actin from Figure 1H. **(J)** Relative mRNA time-course expression of Nipsnap1 (above) and Ucp1 (below) from mice treated with CL 316,243 or PBS IP injection for 1, 4, and 8 days compared to PBS-treated controls (n=5). **(K)** Diagram of Nipsnap1 gene with the mitochondrial targeting sequence indicated (top); Representative Western blot for Nipsnap1 expression in different subcellular compartments from the BAT tissue of 10 day cold-exposed WT mice (bottom). **(L)** Mitochondrial protease protection assay on mitochondria isolated from the BAT tissue of 10 day cold-exposed W7 T mice (bottom). qPCR data are represented as mean ± SEM. Significance is denoted as *p < 0.05, **p < 0.01 ***p < 0.001 by Student’s t test

Nipsnap1 protein induction was then verified by western blot, confirming a significant increase in mice exposed to chronic cold conditions similar to UCP1—the classical marker of thermogenic activation, indicating that the Nipsnap1 expression pattern strongly correlates with other thermogenic proteins (Figure 1E and 1F). The thermogenic properties were specific to only BAT, as there was no increase in inguinal white adipose tissue (iWAT) or in other tissues such as the brain and liver that are reported to have high expression of Nipsnap1 (Fathi, Yarbro and Homayouni, 2021) (Figure S1A-D). Moreover, there was also no increase in Nipsnap2, which shares strong homology with Nipsnap1 (Figure S1E).

To map the expression dynamics of Nipsnap1 in BAT, we measured the protein and mRNA expression of *Nipsnap1* at 1, 4, and 8 days representing acute, intermediate, and prolonged murine cold challenge conditions. Intriguingly, *Nipsnap1* mRNA and protein induction did not begin until day 4 of cold exposure, unlike that of *Ucp1* mRNA and protein levels that are robustly induced after just 1 day of cold exposure (Fig 1G and Figure S1F). This data suggest that Nipsnap1 may play a role in the maintenance rather than the initiation of thermogenesis. Similar induction and expression dynamics of Nipsnap1 was observed in pharmacologically activated thermogenesis using the β_3_ receptor agonist CL 316, 243 (Figure 1H-J), which indicates that Nipsnap1 is regulated, at least in part, through the classical cAMP pathway that induces many other thermogenic genes.

Nipsnap1 contains a mitochondrial-targeting sequence and has been shown to localize to the mitochondrial matrix in brain and liver previously (Tummala, Li and Homayouni, 2010). To reveal the localization of Nipsnap1 in BAT, we fractionated BAT from chronic 10-day cold-exposed wildtype mice and probed cytoplasmic, nuclear, and mitochondrial compartments for Nipsnap1 expression. Indeed, Nipsnap1 was localized to the mitochondrial compartment in BAT (Figure 1K) and is located in the inner mitochondria as shown by mitochondrial protease protection assay (Figure 1L). Taken together, Nipsnap1 is a mitochondrial protein that exhibits a late-stage activation signature in response to cold and β_3_ adrenergic stimuli suggesting that it may play a role in non-shivering thermogenic maintenance rather than activation.

### Mice lacking Nipsnap1 in brown adipose tissue can no longer maintain effective non-shivering thermogenesis

To determine if Nipsnap1 plays a role in the maintenance of non-shivering thermogenesis in BAT, we generated a conditional Nipsnap1 knockout (KO) mouse in thermogenic adipose tissue termed N1-KO or N1-CRE (used interchangeably) (Figure 2A). The Nipsnap1 KO mice are fertile and viable with no obvious developmental defects. We confirmed that Nipsnap1 gene and protein levels were successfully reduced in BAT while remaining unchanged in liver and brain. Importantly, the expression of Nipsnap2, a close homolog of Nipsnap1, was unaffected in N1-KO mice (Figure 2B and 2C). To determine if Nipsnap1 is required for long-term thermogenic maintenance, we placed N1-KO and N1-Flox control mice in metabolic cages to analyze whole-body metabolism. Mice were acclimated in the metabolic cages for 3 days at thermoneutral (TN) temperature conditions (30°C) before subsequent exposure to a 10-day chronic cold challenge. As expected, the shift from TN to cold temperatures resulted in a boost in energy expenditure, but we observed no significant differences between the N1-Flox control and N1-KO mice under TN conditions or in the early stages of cold exposure (thermogenic activation). Remarkably, however, a significant decline in whole-body energy expenditure emerged after 7 days of prolonged cold exposure (the thermogenic maintenance stage), which strikingly aligned with the period of maximum Nipsnap1 expression (day 7-10). Notably, these defects occurred during the night cycle when the mice are the most active (Figure 2D and 2E). The impaired energy expenditure persisted despite no differences in total food consumption and locomotor activity between the N1-KO mice and the N1-Flox controls (Figure S2A and S2B). Significant defects were also observed in VO_2_ and VCO_2_ levels during prolonged cold challenge conditions (Figure 2F and G and Figure S2C and S2D). Furthermore, the N1-KO mice displayed an elevated respiratory exchange ratio revealing a metabolic shift from lipid to carbohydrate metabolism, suggesting a potential defect in the ability to metabolize lipids (Figure 2H and Figure S2E). To determine if the defect in energy expenditure observed during the late-stage thermogenic maintenance period in the N1-KO mice translated to disordered whole-body thermoregulation—a proxy for BAT function, we assessed daily body temperature of N1-KO mice compared to N1-Flox controls under chronic cold conditions for 14 days. Consistent with what was observed previously with energy expenditure, the N1-KO and N1-Flox mice were both able to successfully protect their body temperature against cold in the early stages of thermogenesis. Quite remarkably however, N1-KO mice began to exhibit defects in thermoregulation at day 7 which aligns with the Nipsnap1 maximal expression period (Figure 2I and 2J). Tissue level analysis revealed that the BAT in N1-KO mice displayed a significant decrease in tissue weight while iWAT and gWAT depots were unaffected (Figure 2K and Figure S2F). No differences in overall body weight, fat, or lean mass were observed (Figure S2G-S2I). Taken together, Nipsnap1 ablation in BAT leads to an inability to sustain thermogenic activation. N1-KO mice, when compared to N1-Flox controls, are initially able protect their body temperature when faced with a cold challenge through an active thermogenic process, but later succumb to failures in their ability to maintain effective NST due to their inability to express Nipsnap1.

**Figure 2.**
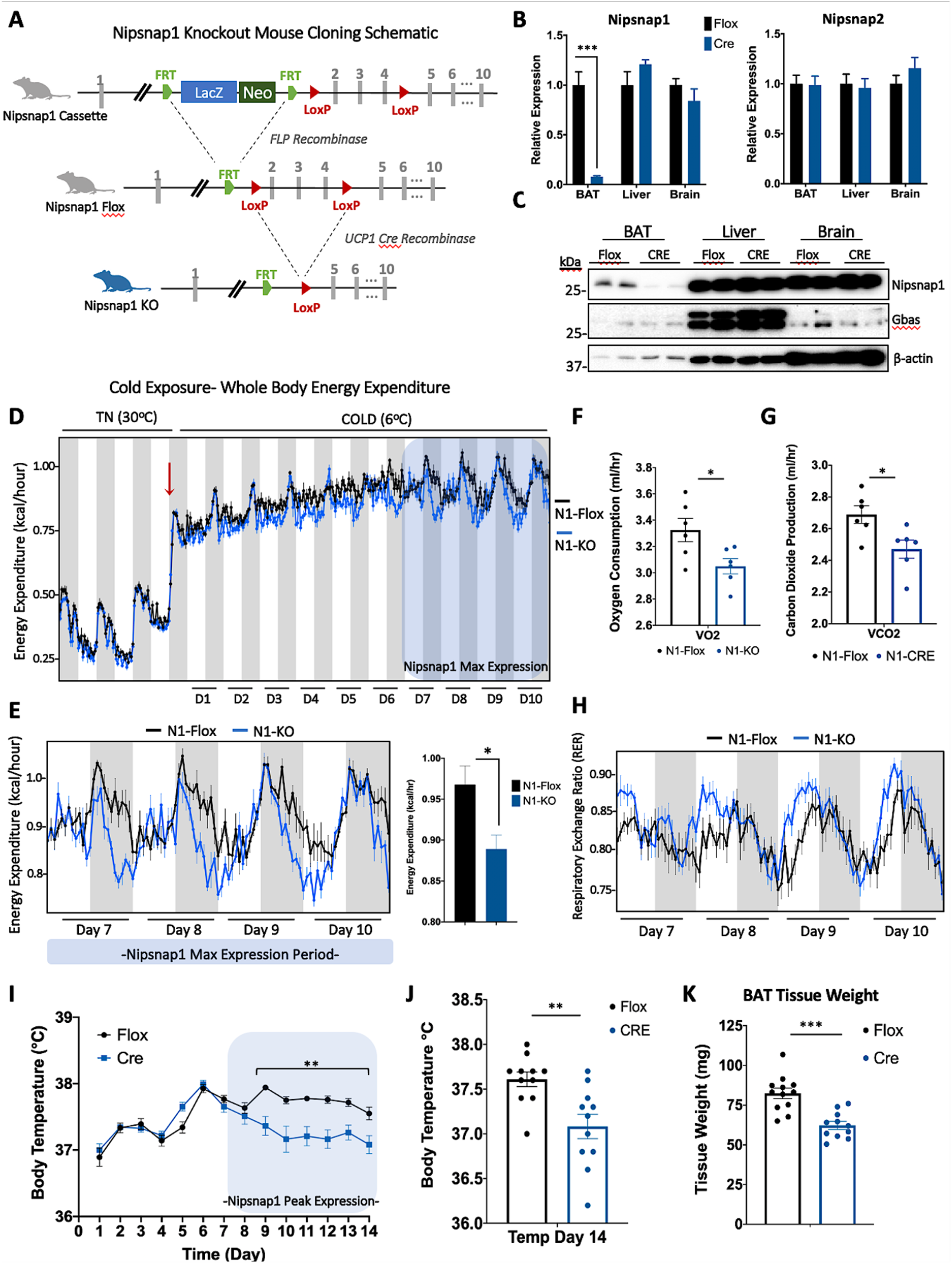
Nipsnap1 ablation in BAT causes severe defects in NST maintenance. Diagram of breeding strategy to generate conditional Nipsnap1 knockout mouse in thermogenic adipose tissue. Relative Nipsnap1 and Nipsnap2 mRNA expression in prolonged cold-induced N1-Flox and N1-KO mice in BAT, Liver, and Brain tissue (n=5). **(C)** Representative Western blot for Nipsnap1 and Nipsnap2 from prolonged cold-induced N1-Flox and N1-KO mice. **(D)** Energy expenditure of N1-Flox and N1-KO mice exposed to thermoneutral (TN, 30°C) for 3 days followed by prolonged cold (COLD, 6°C, red arrow) for 10 days (n=6). Grey bars denote the dark cycle and white bars indicates the light cycle. Blue shaded region indicates an overlay of Nipsnap1 maximal expression period. Red arrow indicates the switch to cold temperatures **(E)** Zoomed-in diagram of energy expenditure plot (from the blue shaded region from 2D) of N1-Flox and N1-KO mice representing day 7-10 of cold exposure (COLD, 6°C). Quantification is indicated to the right. **(F)** Oxygen consumption of N1-Flox and N1-KO (n=6) mice at cold-exposed day 7-10 (n=6). **(G)** Carbon dioxide production of N1-Flox and N1-KO (n=6) mice at cold-exposed day 7-10. **(H)** Respiratory exchange ratio (RER) measurement of N1-Flox and N1-KO (n=6) mice on day 7-10 exposed to cold (COLD, 6°C) **(I)** Daily rectal temperature measurement of N1-Flox and N1-KO exposed to cold (COLD, 6°C) for 14 days (n=11). **(J)** Quantification of figure 2I **(K)** BAT tissue weight from N1-Flox and N1-KO mice exposed to cold for 14 days (n=1101). Whole body metabolic assessments **(D-H)** were analyzed by two-way ANOVA. All other figures unless otherwise indicated are data represented as mean ± SEM. *p < 0.05, **p < 0.01 ***p < 0.001 by Student’s t test.

### Ablation of Nipsnap1 leads to severe defects in BAT lipid metabolism

Given the inability of the Nipsnap1 knockout mice to sustain long-term non-shivering thermogenesis and their corresponding decline in whole-body energy expenditure, we interrogated whether the defects in NST originate specifically from brown fat. We therefore isolated and cultured primary brown adipocytes from N1-Flox and N1-KO mice and assessed cellular respirometry using the Seahorse Bioanalyzer. Indeed, significant defects in cellular respiration were observed in N1-KO adipocytes, impacting basal, CL-induced, and uncoupled respiratory capacity (Figure 3A and B). Based on these results and the known defects in NST maintenance and energy metabolism *in vivo*, we hypothesized that Nipsnap1 deficiency could be compromising classical UCP1-dependent thermogenesis. To test this, we performed thermogenic protein and gene profiling in BAT. Surprisingly, there were no differences in classical thermogenic markers or core mitochondrial genes at either the protein or RNA level (Figure 3C and Figure S3A-S3C). To gain mechanistic insight as to how the ablation of Nipsnap1 could be causing such drastic defects in the maintenance of non-shivering thermogenesis, we performed TMT-labeled proteomics in the brown adipose tissue of our N1-KO mice compared to N1-Flox controls exposed to chronic cold conditions for 10 days to interrogate proteome changes during the thermogenic maintenance phase. We observed a host of significantly differentially expressed proteins and subsequent gene ontology analyses revealed that the majority of the down-regulated proteins were associated with lipid metabolism processes (Figure 3D and 3E). Indeed, many core lipid metabolism proteins such as ATP Citrate Lyase (Acly), Fatty Acid Synthase (Fasn), and Stearoyl CoA Desaturase (Scd1) were significantly reduced in N1-KO mice compared to N1-Flox controls (Figure 3F-3H). Curiously, there were no changes at the mRNA level for lipid metabolism genes, which indicates a potential role for Nipsnap1 in the post-translational regulation of lipid metabolism proteins (Figure S3D). Given these significant changes in lipid metabolism protein levels, we postulated that either lipid accumulation, lipid signaling capacity, and/or lipid beta-oxidation may be impacted. In order to determine if ablation of Nipsnap1 in BAT led to defects in lipid storage capacity, we performed Oil Red O staining in primary brown adipocytes from N1-KO mice compared to controls. Ablation of Nipsnap1 did not alter lipid content or the morphology of the lipid droplet (Figure 3I and Figure S3E). We next examined whether loss-of-Nipsnap1 conferred defects in key lipid proteins in response to norepinephrine (NE) signaling. To test this, we interrogated levels of phosphorylated hormone-sensitive lipase (P-HSL) and phosphorylated acetyl CoA carboxylase (P-ACC) in primary brown adipocytes from N1-KO and N1-Flox mice exposed to NE. As observed in Supplemental Figure 3F, we noted no differences in lipid signaling in response to acute NE treatment. Finally, we wanted to assess whether functional beta-oxidation capacity was affected. We therefore performed cellular respirometry analyses in primary brown adipocytes from N1-KO mice compared to N1-Flox controls in the presence or absence of the carnitine palmitoyl transferase 1 (CPT1) inhibitor etomoxir. As anticipated, there were severe defects in the oxidative capacity of N1-KO primary brown adipocytes compared with N1-Flox controls at basal, oligomycin uncoupled, and maximal respiration (Figure 3J and 3K). The impaired beta-oxidation capacity in primary brown adipocytes prompted us to assess whether these defects in fatty acid metabolism were also present *in vivo*. To test this, we isolated BAT mitochondria from mice that were chronically exposed to a 10-day cold temperature challenge and performed palmitate-driven respirometry analyses in isolated mitochondria from N1-KO mice compared to N1-Flox controls. Remarkably, there was also a significant defect in beta-oxidation capacity present in isolated chronically cold-activated mitochondria from N1-KO mice compared to N1-Flox controls (Figure 3L). This defect in mitochondrial beta-oxidation capacity mechanistically supports our earlier findings in which we observed a macronutrient metabolism shift in N1-KO mice to favor carbohydrate metabolism over the oxidation of lipids due to impaired lipid metabolism (Figure 2H). Taken together, the significant defects in lipid beta-oxidation capacity observed in both isolated mitochondria from BAT tissues, as well as in primary brown adipocytes from N1-KO mice compared to N1-Flox controls mechanistically contributes to the inability to sustain effective non-shivering thermogenesis when faced with a chronic cold challenge. These results suggest that Nipsnap1 integrates with lipid metabolism and when ablated in BAT, leads to significant declines in beta-oxidation capacity.

**Figure 3.**
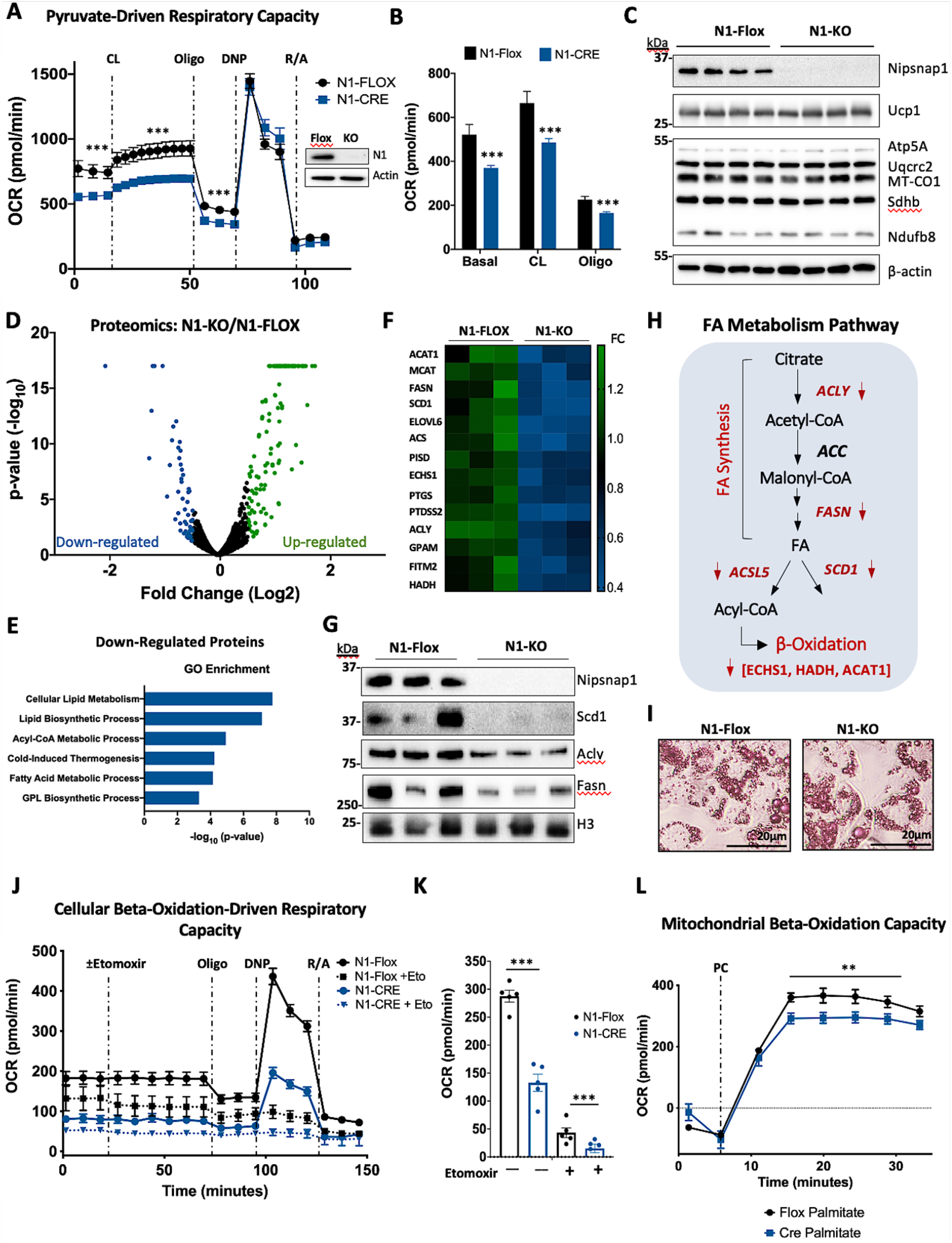
Ablation of Nipsnap1 results in severe defects in BAT Lipid Metabolism. **(A)** Oxygen consumption rate of N1-Flox and N1-KO primary brown adipocytes (n=10). CL, CL 316, 243; Oligo, Oligomycin; DNP, 2,4-Dinitrophenol; R/A, Rotenone and Antimycin A. The western blot inlay represents deletion of Nipsnap1 **(B)** Quantification of Basal, CL 316, 243 and oligomycin-treated oxygen consumption rate of N1-Flox and N1-KO primary brown adipocytes (n=10). **(E)**Representative Western blot of BAT from 10-day cold-induced N1-Flox and N1-KO mice. **(D)** Volcano plot of the proteins upregulated (green) and downregulated (blue) from the BAT tissue proteomic analysis of N1-KO mice compared to N1-Flox controls after 10-day cold exposure. **(E)** Gene ontology analysis of the down-regulated proteins from N1-KO BAT proteome compared to N1-Flox BAT in 10-day cold exposure. **(F)** Heatmap representing the lipid metabolic genes that were downregulated in N1-KO BAT compared to N1-Flox after 10 days of cold exposure. **(G)** Representative Western blot validating select down-regulated proteins involved in lipid metabolism in BAT from N1-KO and N1-Flox mice after 10 days of cold exposure. **(H)** Diagram depicting the down-regulated proteins in the lipid metabolic pathway. **(I)** Representative Oil red O staining image of differentiated N1-Flox and N1-KO primary brown adipocytes. **(J)** Beta-oxidation-driven respiratory capacity with the Seahorse Bioanalyzer in the N1-Flox and N1-KO primary brown adipocytes (n=5). **(K)** Quantification of beta-oxidation-driven basal oxygen consumption rate of N1-Flox and N1-KO primary brown adipocytes (n=5). **(L)** β-oxidation-driven1o4xygen consumption rate of isolated BAT mitochondria from 10-day cold-exposed N1-Flox and N1-KO mice. PC, Palmitoyl-L-carnitine. All Seahorse data are represented as mean ± SEM. *p < 0.05, **p < 0.01 ***p < 0.001 by Student’s t test.

### NIPSNAP1 KO mice do not have increased susceptibility to diet-induced obesity due to enhanced metabolic compensation

To determine if the defect in NST would render N1-KO mice more susceptible to diet-induced obesity, we challenged the mice with 24 weeks of a high fat diet (HFD) feeding regimen. Given that Nipsnap1 is minimally expressed under thermoneutral conditions, we interrogated the response to HFD under cold conditions in which Nipsnap1 would be robustly expressed (Figure S4A and S4B). We first assessed weekly weight accumulation in HFD fed N1-KO mice compared to N1-Flox controls. Both the N1-KO and N1-Flox mice responded to the HFD treatment and gained weight equally throughout the 24-week period (Figure 4A). There were no significant differences in body composition, as assessed by NMR, or in thermogenic tissue weights after HFD feeding (Figure 4B-4C and Figure S4C). To assess if Nipsnap1 played a regulatory role in whole-body glucose metabolism in response to a high fat diet, we performed glucose and insulin tolerance tests (GTT and ITT respectively). There were no differences in GTT or ITT between groups (Figure 4D-E). Taken together, we concluded that although Nipsnap1-KO mice had significant defects in the long-term maintenance of non-shivering thermogenesis, it appeared that this defect did not make the N1-KO mice more susceptible to diet-induced obesity. To interrogate further and given that ablation of Nipsnap1 in the BAT led to significant defects in energy expenditure under chronic cold challenge chow-fed diet conditions (Figure 2D and 2E), we wanted to assess whether this energetic defect persisted under the same chronic cold challenge conditions in the presence of a high fat diet. To test this, we measured whole-body metabolism using the Promethion metabolic cages in N1-KO mice after 24-weeks of a HFD treatment compared to N1-Flox controls under cold environmental conditions. Remarkably, N1-KO mice displayed an increase in energy expenditure over the N1-Flox controls, rather than the decrease they exhibited under chow-fed conditions (Figure 4F). We also observed a corresponding increase in VO_2_ and VCO_2_ levels (Figure S4D and S4E). This increase in energy expenditure was observed most notably in the light cycle when the mice were at rest and was increased independently of changes in food intake or locomotion (Figure 4G and Figure S4F and data not shown). Furthermore, in contrast to the preference observed previously to metabolize carbohydrates instead of fats due to defects in lipid metabolism in the N1-KO mice compared to N1-Flox controls (Figure 2H), we now observed a normalization in the respiratory exchange ratio of the N1-KO mice back to levels of the N1-Flox controls, suggesting a restoration in the ability to metabolize fats as an energy source (Figure S4G). This enhanced energy expenditure and restoration of lipid substrate utilization in N1-KO mice under chronic cold-exposed HFD conditions suggest that the enriched nutrient density and the increased lipid availability in the HFD may be triggering a compensatory mechanism. The increased energy expenditure in N1-KO mice compared to N1-Flox controls could also mechanistically explain why the loss of Nipsnap1 did not render mice more obese than the controls. To determine if the restoration of lipid metabolism was observed at the protein or mRNA level, we profiled select thermogenic and lipid genes. Thermogenic genes and proteins were unaltered as observed previously, but unlike the impaired lipid metabolism signatures that were seen under chronic cold chow-fed diet conditions (Figure 3F and 3G), we no longer noted any significant defects in target lipid metabolism proteins such as Acly and Acsl5 (Figure 4H and Figure S4H-S4J). In summary, the ablation of Nipsnap1 in the BAT which results in the significant decline in NST maintenance capacity under cold chow-fed dietary conditions does not confer increased susceptibility to diet-induced obesity when N1-KO mice are challenged to high fat diet feeding. Instead, due to the increased nutrient density and lipid availability from the high fat diet feeding, the N1-KO mice are able to trigger a compensatory program to increase their energy expenditure and restore lipid substrate utilization to protect themselves from excessive weight gain.

**Figure 4.**
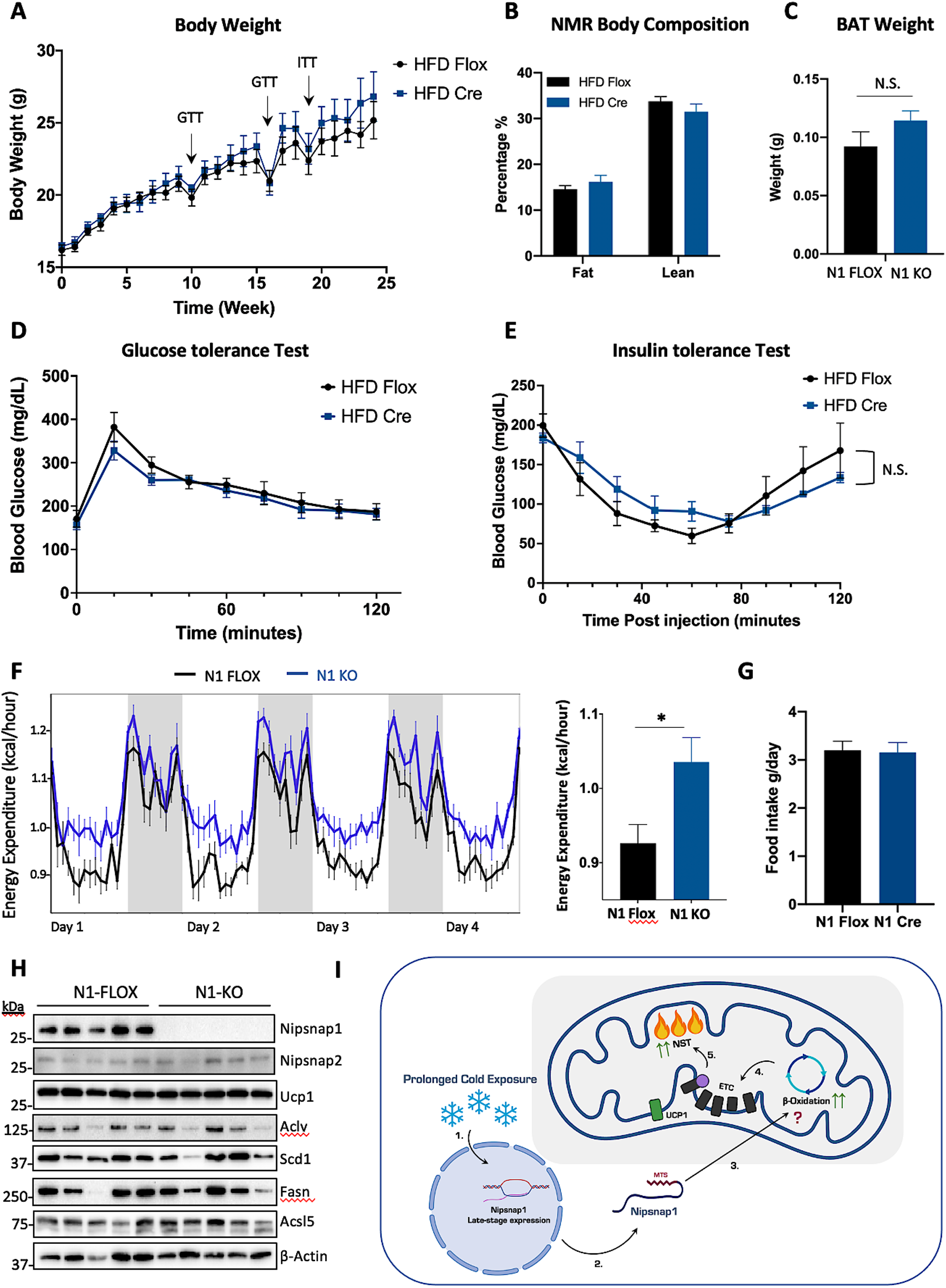
Nipsnap1 KO mice are protected from excessive weight gain through an increase in energy expenditure. All data below are from the N1-Flox and N1-KO mice fed a high fat diet for 24 weeks mice in a chronic cold environment (6oC) (n=5) **(A-C)** Bodyweight measurement, NMR body composition, and BAT tissue weight. GTT, glucose tolerance test; ITT, insulin tolerance test. **(D-E)** Glucose tolerance and insulin tolerance test. **(F)** Energy expenditure. Grey bars indicate dark cycle and white bars indicate light cycle. Quantification is indicated to the right. **(G)** Cumulative daily food intake for one week (wk 23-24). **(H)** Representative Western blot for proteins involved in thermogenesis and lipid metabolism. All data unless otherwise indicated are represented as mean ± SEM. *p < 0.05 by Student’s t test. **(I)** Model for Nipsnap1 function. 1. Chronic cold activation stimulates the expression of Nipsnap1. 2. This results in an accumulation of Nipsnap1 protein levels and transfer into the mitochondria. 3. There Nipsnap1 integrates with lipid metabolism and beta-oxidation processes to 4. ensure sufficient energetic input to both the electron transport chain (ETC). 5. This, along with the action of UCP1 allows for the continued maintenance of the NST program.

## Discussion

Advancing methods to target brown adipose tissue to activate non-shivering thermogenesis and increase energy expenditure has been sought after for decades (Rothwell and Stock, 1979; Warwick and Busby, 1990). To date, the field has made some outstanding advances to identify novel pathways such as succinate metabolism, and futile creatine cycling, in addition to many others that can be targeted to safely activate this protective metabolic process (Kazak *et al*., 2015; Mills *et al*., 2018; Rahbani *et al*., 2021; Sun *et al*., 2021). Despite these advances however, the BAT-mediated increase in energy expenditure in both human and rodent models only persists when BAT is in its activated state (LEAN *et al*., 1988; Deng *et al*., 2018). BAT activation is known to be temporal, so once the activation stimulus, such as cold exposure, is removed, BAT is rapidly deactivated within hours and all protection is lost. The mechanisms surrounding how BAT maintains thermogenic function once initially activated, or the regulatory factors involved with the BAT maintenance process are completely unknown and represents a critical knowledge gap in the field.

Here we have identified Nipsnap1 that displays potent late-stage thermogenic properties in response to both cold environmental exposure and pharmacological β_3_-adrenergic activation. Nipsnap1 exhibits a late-stage thermogenic response that makes it suitable for consideration as a potential thermogenic maintenance factor instead of a novel candidate involved in NST activation. Indeed, the Nipsnap1 KO mice were fully capable of activating the thermogenic program in response to environmental or pharmacological challenges, but then later fail to sustain this activation in the maintenance stage of thermogenesis. Furthermore, the timing of the decline in VO_2_ and VCO_2_ respiratory capacities in the N1-KO mice compared to N1-Flox controls aligned perfectly with the Nipsnap1 expression dynamics. This mechanistically supports the findings that N1-Flox control mice with ample amounts of Nipsnap1 are able to switch seamlessly from thermogenic activation to the thermogenic maintenance phase, whereas the Nipsnap1 KO mice failed to effectively transition to the maintenance stage and were not able to retain the benefit of the initial NST activation. The result was a significant late-stage decline in both energy expenditure and an inability to protect their body temperature when under chronic cold challenge conditions in the N1-KO mice compared to controls. Nipsnap1 did not appear to affect classical thermogenic effector proteins, but rather interfaced with lipid metabolism. Fatty acid beta-oxidation was significantly impaired in N1-KO primary brown fat cultures as well as in *ex vivo* isolated mitochondrial from chronically cold-exposed mice. Upstream lipid metabolism and regulation was unaltered as both lipid storage and signaling capacities were preserved between the KO and the controls.

The manner in which Nipsnap1 regulates late-stage NST and how it integrates with lipid metabolism however is currently unknown and warrants further investigation. Nipsnap1 is evolutionarily conserved which argues a critical role for the protein although the specific function as well its protein functional domains have yet to be identified (Seroussi *et al*., 1998). Reports have indicated that Nipsnap1 can be localized to either the mitochondrial surface (Yamamoto, S. *et al*., 2017) or the mitochondrial matrix (Nautiyal, M. *et al*, 2010), or both in response to mitochondrial damage (Abudu, Y.P. *et al*., 2019). In this present study, we demonstrate that Nipsnap1 is localized to the mitochondrial matrix in BAT. It is therefore mechanistically plausible that Nipsnap1 could both directly bind and/or directly regulate the function of proteins involved in lipid beta-oxidation. Further experiments are warranted to tease apart the detailed mechanistic interface of the Nipsnap1-lipid metabolism axis.

## ACKNOWLEDGMENTS

We acknowledge the Cornell Biotechnology Resource center for the mitoproteomics and whole tissue proteomic analysis. We also sincerely thank E.D.R, L.K, P.C, P.S, K.S, and B.C. for their critical review of the manuscript. A special thanks to R.B. for the figure graphics.

## AUTHOR CONTRIBUTIONS

Conceptualization, J.J.B. and Y.L.; Methodology, J.J.B., Y.L., and Y.Q.; Investigation, Y.L., Y.Q., C.C., P.C., K.E., and S.X.; Validation, Y.L., S.P., S.E., and N.S. Formal Analysis, J.J.B. and Y.L.; Writing – Original Draft, J.J.B., Y.L., and Y.Q.; Writing – Review & Editing, J.J.B., Y.L., Y.Q., C.C., P.C., K.E., and S.X.; Visualization, J.J.B. and Y.L.; Supervision, J.J.B.

## Declaration of interests

The authors declare no competing interests.

### STAR METHODS

## RESOURCE AVAILABILITY

### Lead contact

Further information and requests for resources and reagents should be directed to and will be fulfilled by the lead contact, Joeva J Barrow

### Materials availability

Mouse lines are available from the Lead Contact upon request.

### Data and code availability

Mitochondrial and whole tissue BAT proteomics data have been deposited at NCBI GEO and will be made publicly available as of the date of publication. Accession numbers are listed in the key resources table. Any additional information required to reanalyze the data reported in this paper are available from the lead contact upon request.

## Key Resources Table

| REAGENT or RESOURCE | SOURCE | IDENTIFIER |
| --- | --- | --- |
| **Antibodies** |  |  |
| Rabbit polyclonal anti-GBAS | Abcam | Cat# ab83845 |
| Rabbit monoclonal anti-UCP1 | Abcam | Cat# ab209483 |
| Total OXPHOS Rodent WB Antibody Cocktail | Abcam | Cat# ms604 |
| Mouse monoclonal anti-MTCO1 | Abcam | Cat# ab14705 |
| Rabbit monoclonal anti-β-Actin | Cell Signaling | Cat# 4970 |
| Rabbit monoclonal anti-NIPSNAP1 | Cell Signaling | Cat# 13226 |
| Rabbit monoclonal anti-GAPDH | Cell Signaling | Cat# 2118 |
| Rabbit polyclonal anti-Histone H3 | Cell Signaling | Cat# 9715 |
| Rabbit monoclonal anti-Tomm20 | Cell Signaling | Cat# 42406 |
| Mouse monoclonal anti-VDAC1 | Santa Cruz | Cat# sc-390996 |
| Rabbit polyclonal anti-FASN | Cell Signaling | Cat# 3189 |
| Rabbit monoclonal anti-SCD1 | Cell Signaling | Cat# 2794 |
| Rabbit polyclonal anti-ATP-Citrate Lyase | Cell Signaling | Cat# 4332 |
| Mouse monoclonal anti-ACSL5 | Santa Cruz | Cat# sc-365478 |
| Rabbit polyclonal anti-HSL | Cell Signaling | Cat# 4107 |
| Rabbit polyclonal anti-Phospho-HSL (Ser563) | Cell Signaling | Cat# 4139 |
| Rabbit polyclonal anti-ACC | Cell Signaling | Cat# 3662 |
| Rabbit polyclonal anti-Phospho-ACC (Ser79) | Cell Signaling | Cat# 3661 |
| **Bacterial and virus strains** |  |  |
| Ad5CMVeGFP | UI Viral Vector Core | N/A |
| Ad5CMVCre | UI Viral Vector Core | N/A |
| **Chemicals, peptides, and recombinant proteins** |  |  |
| Insulin human | Sigma-Aldrich | I2643;<br>CAS: 11061-68-0 |
| Insulin from bovine pancreas | Sigma-Aldrich | I5500;<br>CAS: 11070-73-8 |
| Dispase II | Roche | 04942078001 |
| Collagenase B | Roche | 11088815001 |
| Collagenase D | Roche | 11088866001 |
| Bovine Serum Albumin | Sigma-Aldrich | A7906<br>CAS: 9048-46-8 |
| Bovine Serum Albumin, fatty acid free | Sigma-Aldrich | A6003;<br>CAS: 9048-46-8 |
| Rosiglitazone | Cayman | 71740;<br>CAS: 122320-73-4 |
| Dexamethasone | Sigma-Aldrich | D4902;<br>CAS: 50-02-2 |
| 3-Isobutyl-1-methylxanthine | Sigma-Aldrich | I7018;<br>CAS: 28822-58-4 |
| 3,3',5-Triiodo-L-thyronine sodium salt | Sigma-Aldrich | T6397;<br>CAS: 55-06-1 |
| CL 316,243 (sodium salt) | Cayman | 17499;<br>CAS: 138908-40-4 |
| Oligomycin | Sigma-Aldrich | 495455;<br>CAS: 1404-19-9 |
| 2,4-Dinitrophenol | TCI | D0109;<br>CAS: 51-28-5 |
| Rotenone | Thermo-Fisher | ICN15015401;<br>CAS: 83-79-4 |
| Antimycin A | Sigma-Aldrich | A8674;<br>CAS: 1397-94-0 |
| Sodium Pyruvate (100 mM) | Thermo-Fisher | 11360070 |
| L(-)-Malic acid | Grayline Medical | CAS: 97-67-6 |
| Palmitoyl-L-carnitine chloride | Sigma-Aldrich | P1645;<br>CAS: 18877-64-0 |
| Guanosine 5'-diphosphate sodium salt | Sigma-Aldrich | G7127;<br>CAS: 43139-22-6 |
| FCCP | Cayman | 15218;<br>CAS: 370-86-5 |
| Trypsin | Sigma-Aldrich | T4799;<br>CAS: 9002-07-7 |
| **Commercial assays** |  |  |
| Qproteome Mitochondria Isolation Kit | Qiagen | CAS: 37612 |
| **Oligonucleotides** |  |  |
| See Table S1 | This paper | N/A |
| **Software and algorithms** |  |  |
| AlphaView 3.5.0 | Protein Simple | N/A |
| GraphPad Prism 8 | GraphPad Software | N/A |
| Agilent Wave software v2.4 | Agilent Technologies | N/A |
| SoftMax Pro 6 | Molecular Devices | N/A |
| OLYMPUS cellSens Entry 3.2 | OLYMPUS | N/A |
| **Other** |  |  |
| Blood Glucose Meter | OneTouch | Model# Geeloa |
| Blood Glucose Test Strips |  |  |
| Cell strainer, 100µM | VWR | CAS: 76327-102 |
| Cell strainer, 40µM | VWR | CAS: 76327-098 |
| Promethion metabolic cages | Sable Systems International | N/A |
| Rodent Diet With 60% kcal% Fat | RESEARCH DIETS | CAS: D124921 |
| TissueLyser II | Qiagen | CAS: 85300 |

## EXPERIMENTAL MODEL AND SUBJECT DETAILS

### Mice

All animal experiments were approved by the Institutional Animal Care and Use Committee (IACUC) at Cornell University. To generate the conditional Nipsnap1 transgenic mice, genetic crossing procedures were performed as described previously (Skarnes *et al*., 2011). Briefly, murine heterozygous Nipsnap1 sperm (EMMA Strain ID: EM 04250) harboring loxP sites between exons 2-4 of the Nipsnap1 gene were acquired from the European Conditional Mouse Mutagenesis Program (EUCOMM). *In vitro* fertilization (IVF) with Nipsnap1^Flox/wt^ sperm and WT C57BL/6J mouse eggs was performed at the Cornell Stem Cell and Transgenic Core Facility. Live heterozygous Nipsnap1^Flox/wt^ mice were subsequently crossed with mice harboring the FLP recombinase to remove the Neomycin cassette and inter-crossed to generate the homozygous floxed Nipsnap1^Flox/Flox^ strain. To achieve thermogenic adipose tissue-specific deletion of Nipsnap1, Nipsnap1^Flox/Flox^ mice were then crossed with UCP1 Cre mice All mice were maintained on 12 h light and dark cycles and fed *ad libitum* either standard irradiated rodent chow or a 60% high fat diet (HFD) (Research Diets D12492). For cold exposure experiments, 6-week-old male and female wild-type C57BL/6J (Jackson #000664) and Nipsnap1 transgenic mice were single-housed in rodent incubators and exposed to a constant temperature of 6.5°C for indicated time points. For pharmacological induced thermogenesis experiments, male and female mice at 6 weeks of age were injected daily via intraperitoneal injection (IP) with 1mg/kg CL 316,243 for the indicated time points. HFD treatment was initiated in male and female mice at 6 weeks of age for 24 weeks.

### Mouse Primary Brown Adipocytes Culture and Differentiation

For *in vitro* studies of thermogenesis with primary brown adipocytes, BAT adipose tissue was obtained from 3-week-old male wild-type C57BL/6J mice or Nipsnap1 Flox mice. Fat pads from 5 mice were pooled after dissection, minced thoroughly and digested in 15ml of BAT dissociation buffer (123mM NaCl, 5mM KCl, 1.3mM CaCl_2_, 5.0mM Glucose, 100mM HEPES, 4% BSA and 1.5mg/mL collagenase B) for 30min at 37°C with constant shaking. The cell suspension was filtered with a 100μm cell strainer and centrifuged at 600g for 5min. The pellet was then suspended in adipocyte culture medium (DMEM/F12 with 10% FBS, 25mM HEPES, 1% PenStrep), filtered with a 40μM cell strainer, centrifuged at 600g for 5min, and resuspended in culture medium and plated in 10cm polystyrene cell culture dishes. Preadipocytes were seeded to post confluency and differentiated with DMEM/F12 supplemented with 5ug/mL Insulin, 1μM Rosiglitazone, 1μM Dexamethasone, 0.5mM Isobutylmethylxanthine and 1nM T3. Cells were maintained in differentiation media for 48h before being switched into maintenance media (5ug/mL Insulin and 1μM Rosiglitazone and 1nM T3) for 4-6 days. For primary brown adipocyte adenoviral-GFP or Cre treatment, primary cells were treated with 1000 Multiplicity of infection (MOI) of either Adeno GFP or Adeno Cre virus for 72hours. Cells were then trypsinized and reseeded into desired assay plates and exposed to another dose of adenovirus treatment at an MOI of 1000 for 72h. Cells were then differentiated as described above. For lipid signaling experiments, primary adipocytes were exposed to 1μm of norepinephrine for 1 hour.

## METHOD DETAILS

### Glucose and Insulin Tolerance Tests

Glucose tolerance tests were performed at weeks 10 and 16 of the HFD treatment. Mice were fasted for 16h before receiving 1.5mg/kg intraperitoneal injection (IP) of glucose. Blood glucose levels were measured using a glucometer every 15min for a duration of 2 hours after glucose administration. Insulin tolerance tests were performed at week 19 of the HFD treatment. Mice were fasted for 6h prior to receiving 1U/kg IP injection of insulin. Blood glucose levels were then measured by glucometer every 15min for a period of 2 hours post-injection. The blood sample was collected from tail nicking.

### In Vivo Indirect Calorimetry

Mice were single-housed in rodent Promethion metabolic cages (Sable Systems International) situated in a temperature-controlled thermal cabinet with a 12-hour light/dark cycle schedule. For cold exposure studies, mice were acclimated in metabolic cages set to 30°C for 72 hours after which the thermal cabinet was cooled to 6.5°C for the duration of the cold exposure study. For HFD studies, mice were acclimated in the metabolic cages for 48 hours prior to whole-body metabolic measurements at either 30°C or 6.5°C. Comprehensive real-time metabolic measurements such as energy expenditure (kcal/hr), oxygen consumption (VO_2_), carbon dioxide expiration (VCO_2_), locomotive movement (measured by X-Y infrared beam breaks), and respiratory exchange ratio (RER) were measured and recorded every 3 mins using the Sable System data acquisition software (IM-3 v.20.0.3). Raw data was then processed using the Sable System Macro Interpreter software and One-Click Macro systems (v2.37). Data was further processed using CalR (Mina *et al*., 2018).

### RNA and Protein Analyses

Total RNA was extracted using TRIzol reagent. Isolated tissues were homogenized using the Qiagen TissueLyser II in TRIzol to isolate total RNA and processed according to the manufacturer’s instruction. Total RNA (1ug) was reverse transcribed to cDNA using the qScript cDNA Synthesis kit. RT-qPCR was performed using the CFX384 Real-Time PCR System using SYBR Green. For protein analyses, tissues were homogenized with the Qiagen TissueLyser II in 2% SDS lysis buffer supplemented with protease and phosphatase inhibitors. Protein concentration was measured by bicinchoninic protein assay. Proteins were then resolved on 12% SDS-PAGE. The protein gels were then subjected to sliver staining or were transferred to polyvinylidene fluoride (PVDF) membranes. Blots were probed with target antibodies and visualized using the FluorChem imaging system. Images were quantified with densitometry using the AlphaView software.

### Mitochondrial Isolation

To isolate mitochondria for respirometry analyses, brown fat pads were isolated and mechanically disrupted with scissors in a mitochondrial isolation buffer (300mM sucrose, 5mM HEPES, 1mM EDTA, pH 7.2 with KOH). Minced brown adipose tissue was then lysed using a glass-Teflon homogenizer with a tight-fitting pestle using 15 strokes. Homogenates were filtered by 100 μM mesh filter and centrifuged at 600*g* to remove the nuclei and cellular debris. The supernatant was retained and centrifuged at 8,500*g* for 10 min to isolate mitochondria. Mitochondria were then resuspended in MAS buffer (70mM sucrose, 220mM mannitol, 5mM KH2PO4, 5mM MgCl, 2mM HEPES, 1mM EGTA, pH7.2 with KOH) and quantified by bicinchoninic protein assay. To isolate mitochondria for WB analyses, the Qproteome Mitochondria Isolation Kit was used. For mitochondria protection assays, isolated and purified mitochondria were treated with or without trypsin (150ug/ml) for 30min at RT.

### Cellular and Mitochondrial Respiration Assay

Oxygen consumption rate (OCR) was measured by Seahorse XFe24 analyzer (Agilent). For cellular respirometry analyses, primary brown adipocytes were differentiated in XFe24 cell culture plates as previously described. For cellular CL-induced mitochondrial stress tests, on day 7 post-differentiation, cells were washed and switched to unbuffered DMEM supplemented with 4.5g/L glucose, 4mM glutamine, 100mM pyruvate, and 2% of fatty acid-free BSA, pH7.4 by NaOH. For cellular β-oxidation test, cells were washed and switched to unbuffered DMEM supplemented with 0.5mM glucose, 1mM glutamine, 0.5mM L-carnitine, 1% FBS for starvation for 24h. At the day of the assay, cells were switched into unbuffered DMEM supplemented with 2mM glucose, 0.5mM L-carnitine, incubated for 30min and then put into the assay. The compounds concentrations were as follows (final concentration): For standard mitochondrial stress test, A: CL 316, 243 (10 μM), B: oligomycin (4.5μM), C: DNP (0.6mM), D: Rotenone/Antimycin A (4μM each); for β-oxidation test, A: Etomoxir (10μM). B: Oligomycin (4.5μM), C: DNP (0.6mM), D: Rotenone/Antimycin A (4μM each). For mitochondrial respirometry analyses, fresh mitochondria from interscapular brown adipose tissue were isolated as described previously. Then 5μg of mitochondria were resuspended in 50μl of MAS buffer per well in a seahorse plate. The seahorse plate was centrifuged at 2000g for 20min to adhere the mitochondria. Then 450μl of MAS buffer was carefully added to each well. The mitochondrial stress test compounds were as follows (final concentration): A: Pyruvate (10mM) and Malic acid (5mM) (for pyruvate-driven respiration), Palmitoyl-L-carnitine (50μM) (for palmitate-driven respiration), B: GDP (1mM), C: FCCP (4μM), D: Rotenone/Antimycin A (2μM each). Respirometry data were collected using the Agilent Wave software v2.4.

### Body Composition

Body composition to assess lean and fat mass were measured via NMR using the Minispec LF65 Body Composition Mice Analyzer (Bruker, Karlsruhe, Germany)

### Mitochondrial and Whole Cell Proteomics

To analyze the mitochondrial proteome, total mitochondria were isolated from brown fat tissue as described above. BAT mitochondria were lysed in radio immunoprecipiation buffer (RIPA) [150mM NaCl, 50mM Tris-Cl pH 7.5, 0.5% NP-40, 0.1% Sodium deoxycholate, supplemented with protease and phosphatase inhibitors]. To analyze the whole cell proteome, brown adipose fat pads were isolated and lysed in 2% SDS solution supplemented with protease and phosphatase inhibitors. Sample lysates were quantified by bicinchoninic protein assay and delivered to the Biotechnology Resource Center (BRC) at Cornell University for Tandem Mass-Tagged (TMT) shotgun-based quantitative proteomics. Briefly, proteins were denatured, reduced, cysteine blocked, and digested using the S-trap approach. The resulting trypic peptides were TMT-labeled, and pooled. The labeled peptides were then fractionated by high pH reverse phase chromatography by the Ultimate 3000 MDLC platform into 10 fractions. Then samples were subjected to nanoLC-MS/MS analysis using a reverse phase HPLC separation and NanoLC RP coupled with an Orbitrap Eclipse mass spectrometer (Thermo Scientific) equipped with a nano ion source. Ion quantification and proteomic database searches were conducted using the Proteome Discoverer 2.4 software against the mouse database. All MS and MS/MS raw spectra were processed using Proteome Discoverer 2.3 (PD 2.3, Thermo) for reporter ion quantitation analysis. Gene ontology was performed by EnrichR (E. Y. Chen *et al*., 2013; Kuleshov *et al*., 2016).

### Oil Red O Staining

Oil red O staining was performed as described previously (Kraus *et al*., 2016). In short, Oil Red O stock solution was prepared by dissolving 0.2g Oil red O powder in 40ml isopropanol. A 2:3 stock to distilled water working solution was then prepared and filtered immediately before use. For cell staining, cells were washed with PBS and fixed in 4% formaldehyde in PBS for 15min at RT. Fixed cells were then washed 3 times with PBS, and Oil red O working solution was then added. After 30min incation with mild shaking at RT, cells were washed 5 times in PBS and then imaged with an Olympus IX71 Inverted Fluorescence Microscope. Images were acquired and analyzed by OLYMPUS cellSens Entry 3.2. For staining quantification, 1ml isopropanol was added to the stained cells. The plate was incubated for 10min at RT with mild shaking. 200μl of the eluate per sample was then transferred to a clear bottom 96-well plate along with 200μl pure isopropanol as a blank control. Absorption was measured at 510nm by SpectraMax M3 and data was acquired and interpreted by SoftMax Pro 6.

### QUANTIFICATION AND STATISTICAL ANALYSIS

Statistical analyses were calculated using GraphPad Prism 8 software. Specific statistical tests are indicated in each figure and data are all represented as the mean ± S.E.M. Both unpaired two-tailed Student’s T-test and two-way ANOVA were used. All statistical analyses were performed in consultation with the Cornell Statistical Consulting Unit (CSCU). Power analyses were performed for *in vivo* and *ex vivo* animal experiments. For *ex vivo* cellular experiments, an n=3 at an effect size of > 50% difference between control and treatment groups was determined to yield significance. At least three biological replicates were conducted for all *ex vivo* studies with independently derived primary cells. For animal experiments, an n=5-10 per group for an effect size of >10-20% respectively was determined to yield significance. A minimum of 2 biological replicates were performed for all *in vivo* experiments with appropriate vehicle controls.

**Table 1.**
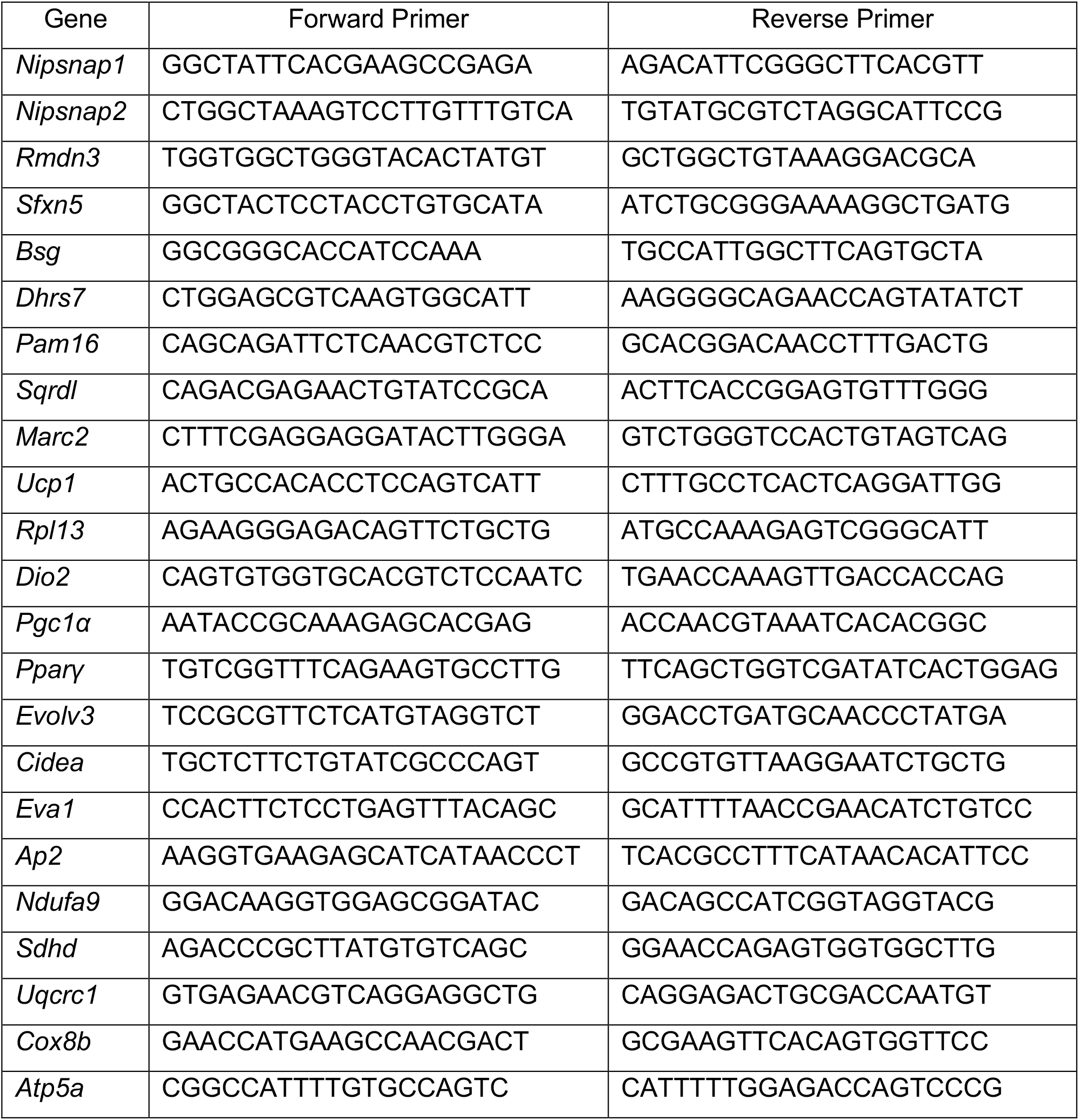
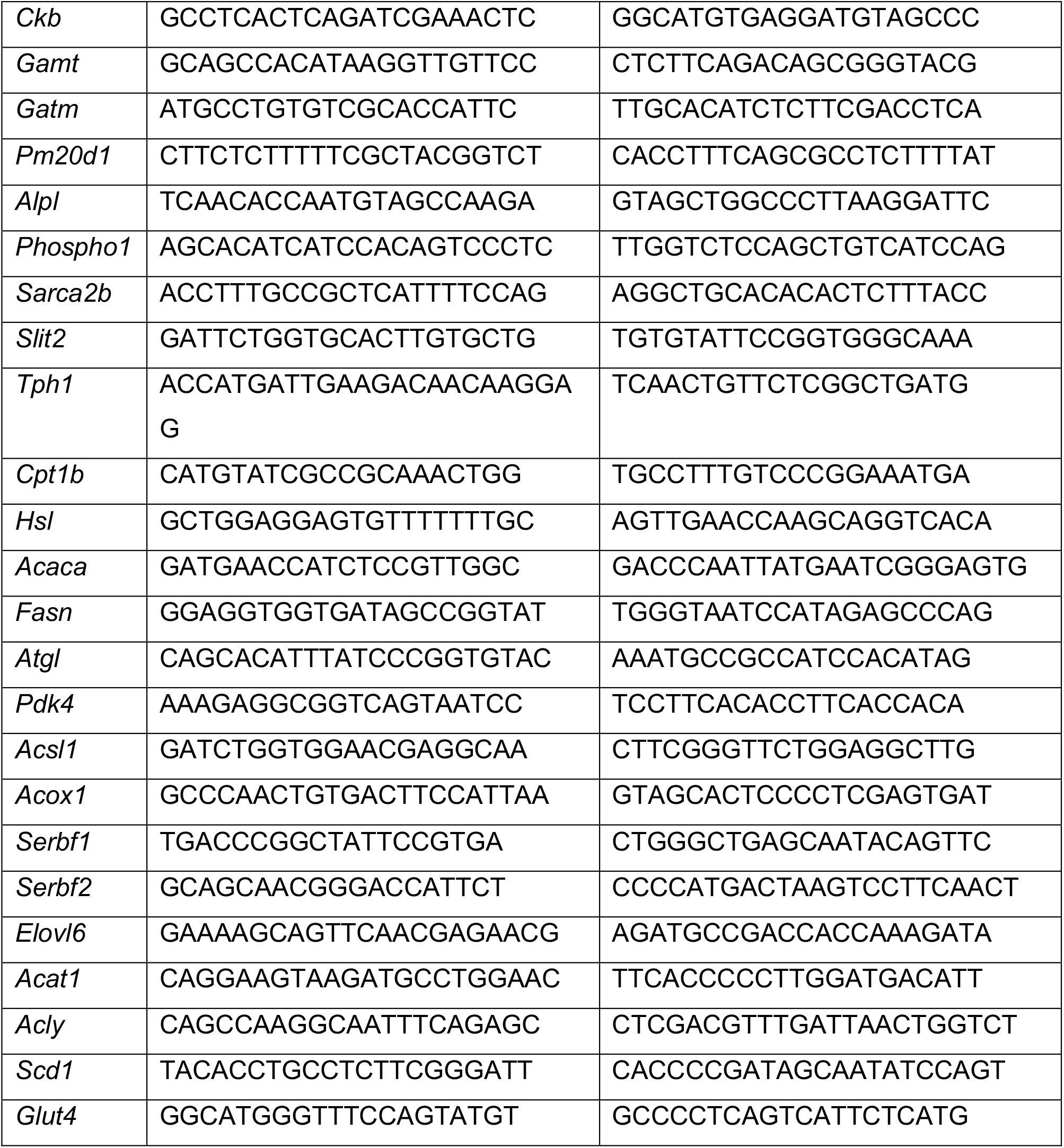

## SUPPLEMENTAL INFORMATION

**Figure S1.**
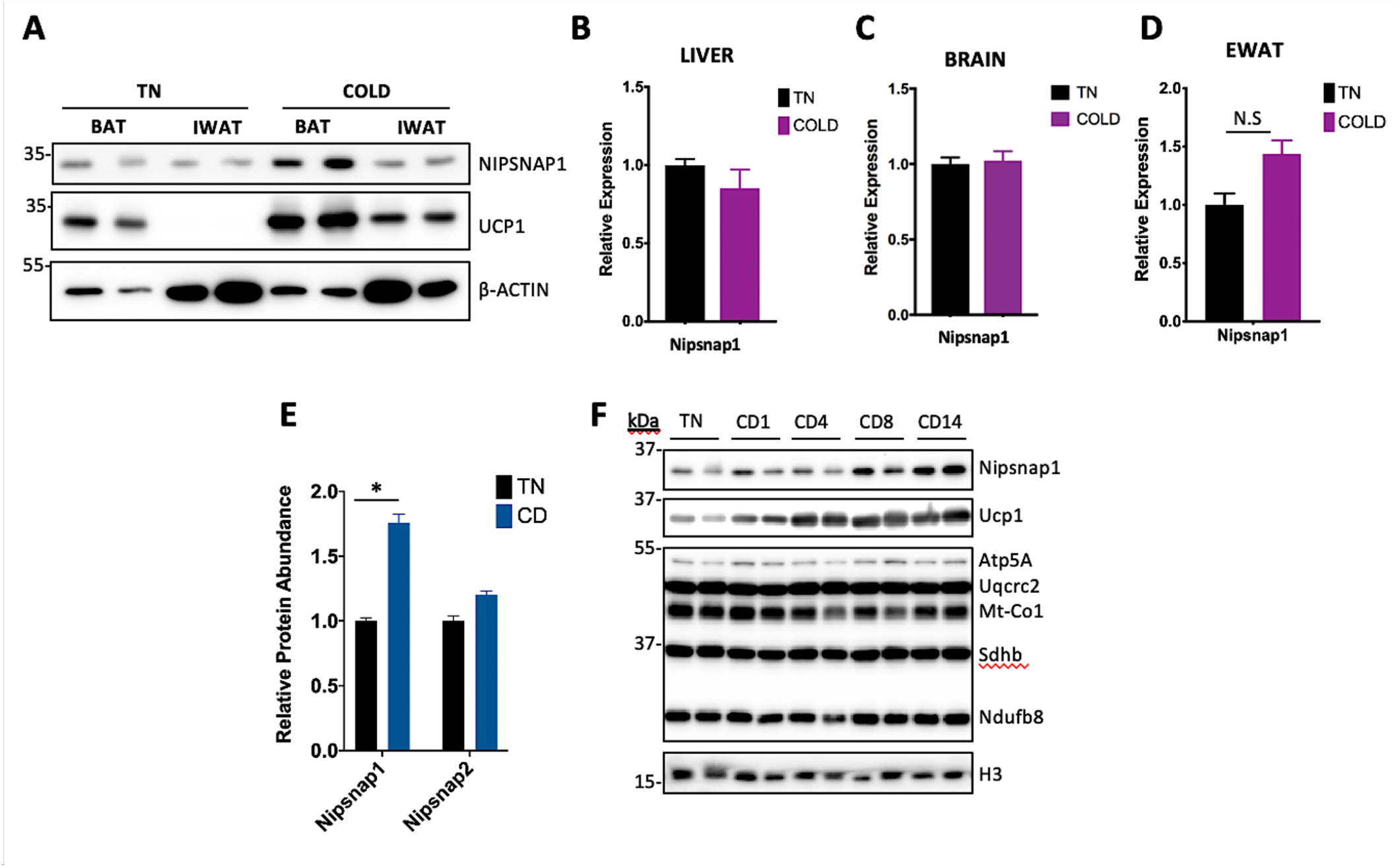
Increases in Nipsnap1 levels in response to cold is BAT-specific (Related to Figure 1) **(A)**Representative Western blot from murine BAT and iWAT tissue in TN and Cold for 8 days. **(B-D)** Relative mRNA expression of Nipsnap1 in murine liver, brain and EWAT tissue exposed to cold for 8 days compared to thermoneutral environment (n=5). **(E)**Protein abundance levels of Nipsnap1 and Nipsnap2 in mice TN and Cold exposed BAT mitochondria proteomic analysis (n=3). **(F)**. Representative Western blot from murine BAT tissue exposed to cold for 1, 4, 8 and 14 days. All mRNA profiling was conducted by qPCR with data represented as mean ± SEM. *p < 0.05 by Student’s t test.

**Figure S2.**
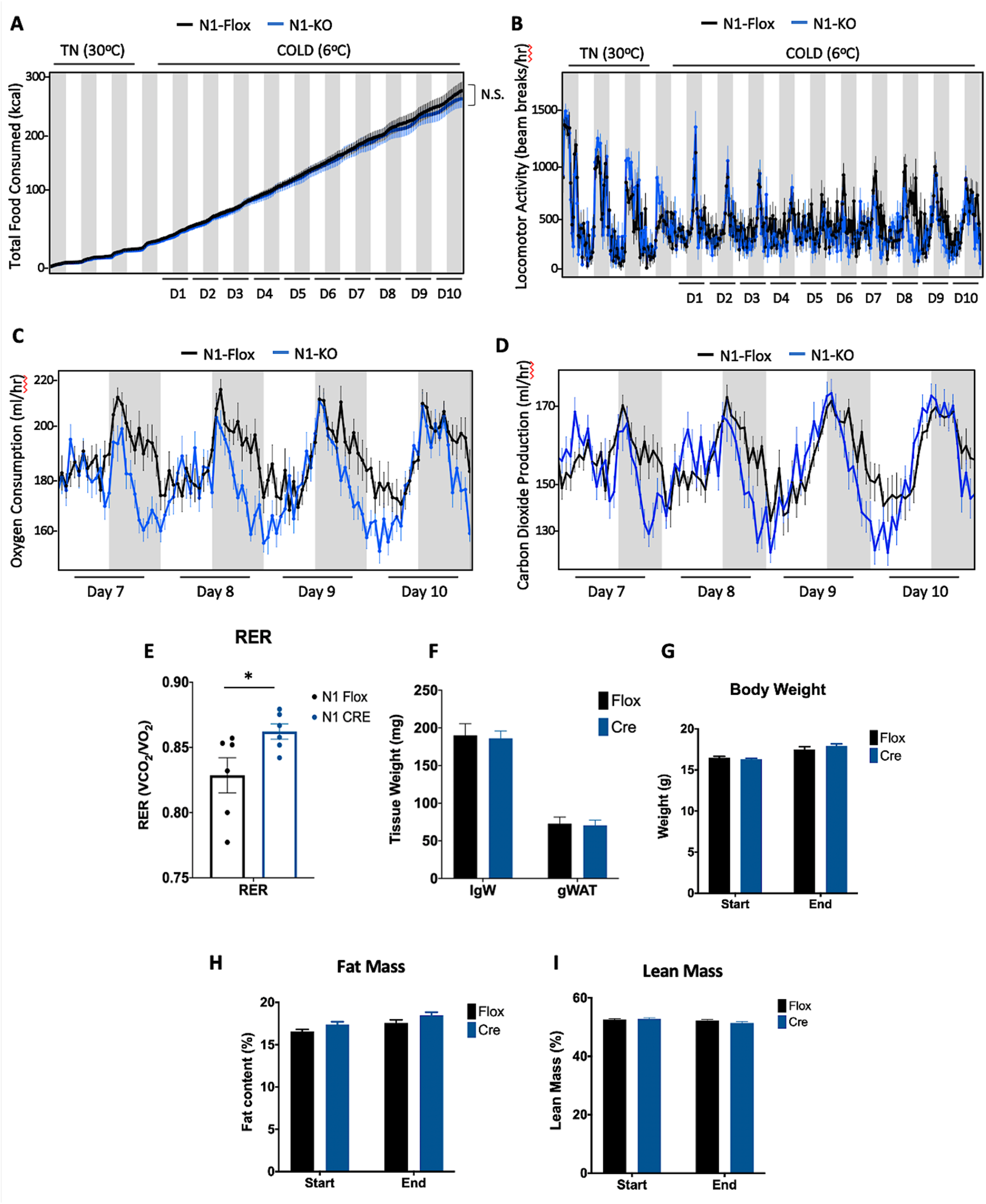
N1-KO mice exhibit defects in VO_2_ and VCO_2_ levels in response to chronic cold (Related to Figure 2) **(A-B)** Total food consumption **(A)** and locomotor activity **(B)** of N1-Flox and N1-KO (n=6) mice exposed at thermoneutral (TN, 30°C) temperature for 3 days followed by prolonged cold exposure (COLD, 6°C) for 10 days. **(C-D)** Oxygen consumption **(C)** and CO_2_ production **(D)** in N1-Flox and N1-KO (n=6) mice exposed to cold during the Nipsnap1 maximal expression period (day 7-10). **(E)** Quantification of the respiratory exchange ratio (RER) in N1-Flox and N1-KO (n=6) mice exposed to cold during the Nipsnap1 maximal expression period (day 7-10). **(F-I)** Tissue weight **(F)**, whole body weight (G), as well as fat **(H)** and lean mass **(I)** content measurement of N1-Flox and N1-KO (n=6) mice exposed at thermoneutral (TN, 30°C) for 3 days followed by prolonged cold (COLD, 6°C) for 10 days. Mouse food intake and locomotor measurement data were analyzed by two-way ANOVA **(A-B)**; All other data are represented as mean ± SEM. *p < 0.05 by Student’s t test. 35

**Figure S3.**
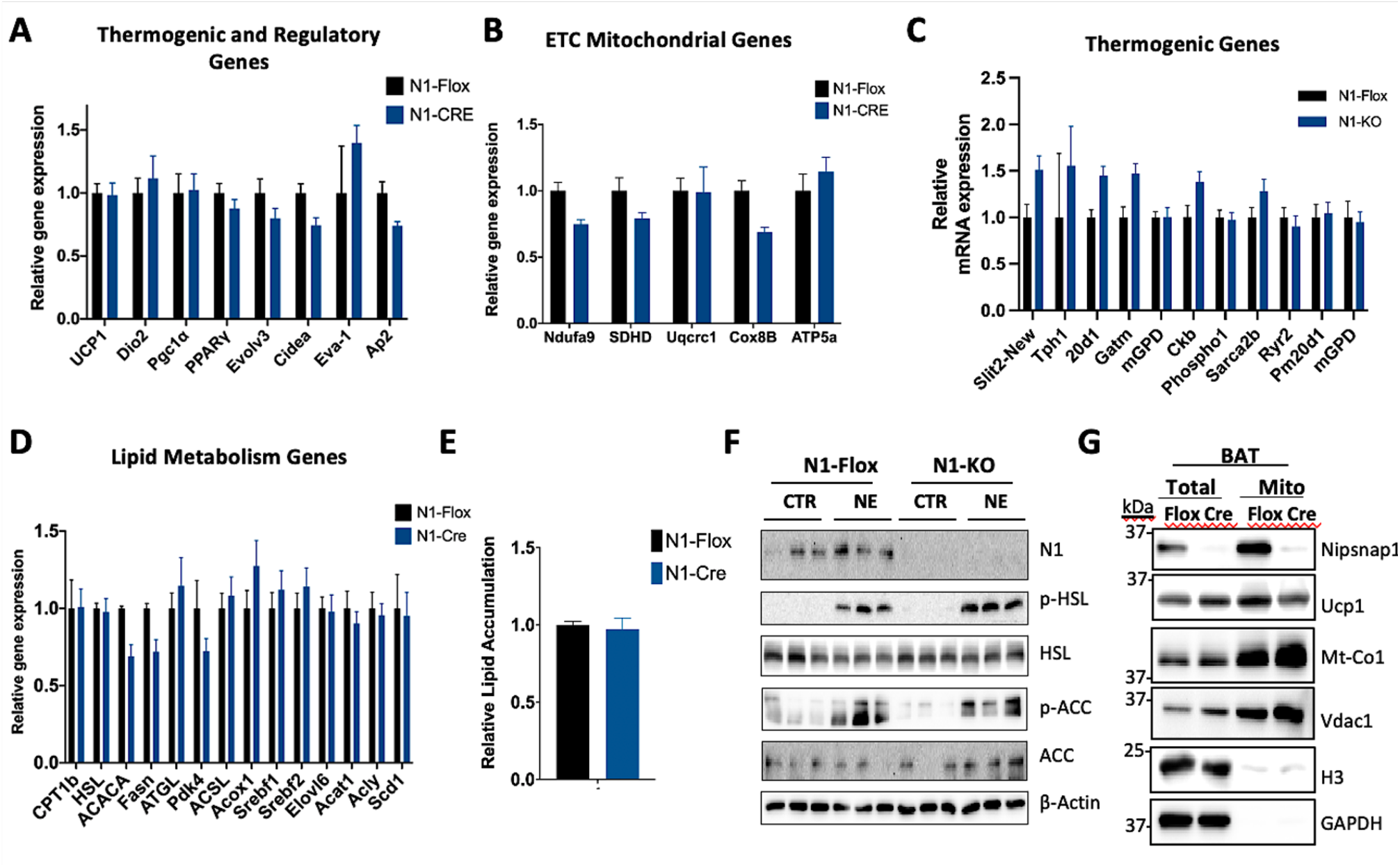
BAT-specific ablation of Nipsnap1 does not affect thermogenic of lipid metabolism genes. (Related to Figure 3) **(A-D)** Relative mRNA expression of thermogenic effector **(A and C)**, mitochondrial **(B)** and lipid metabolism **(D)** genes in BAT tissue from N N1-KO mice compared to N1-Flox mice exposed to 10 days of cold exposure (n=6). **(E)** Quantification of Oil red O staining in differentiated N1-KO compared to N1-Flox primary adipocytes (n=3). **(F)** Representative Western blot for lipid metabolic signaling proteins from primary differentiated N1-KO adipocytes compared to N1-Flox controls treated with 1 μM of norepinephrine (NE) for 1 hr (n=3). **(G)** Representative western blot of isolated mitochondrial from BAT from chronic 10-day exposed N1-Flox and N1-KO mice. All mRNA profiling was conducted by qPCR. mRNA quantification and lipid accumulation measurements are data represented as mean ± SEM. Statistics were performed by Student’s t test.

**Figure S4.**
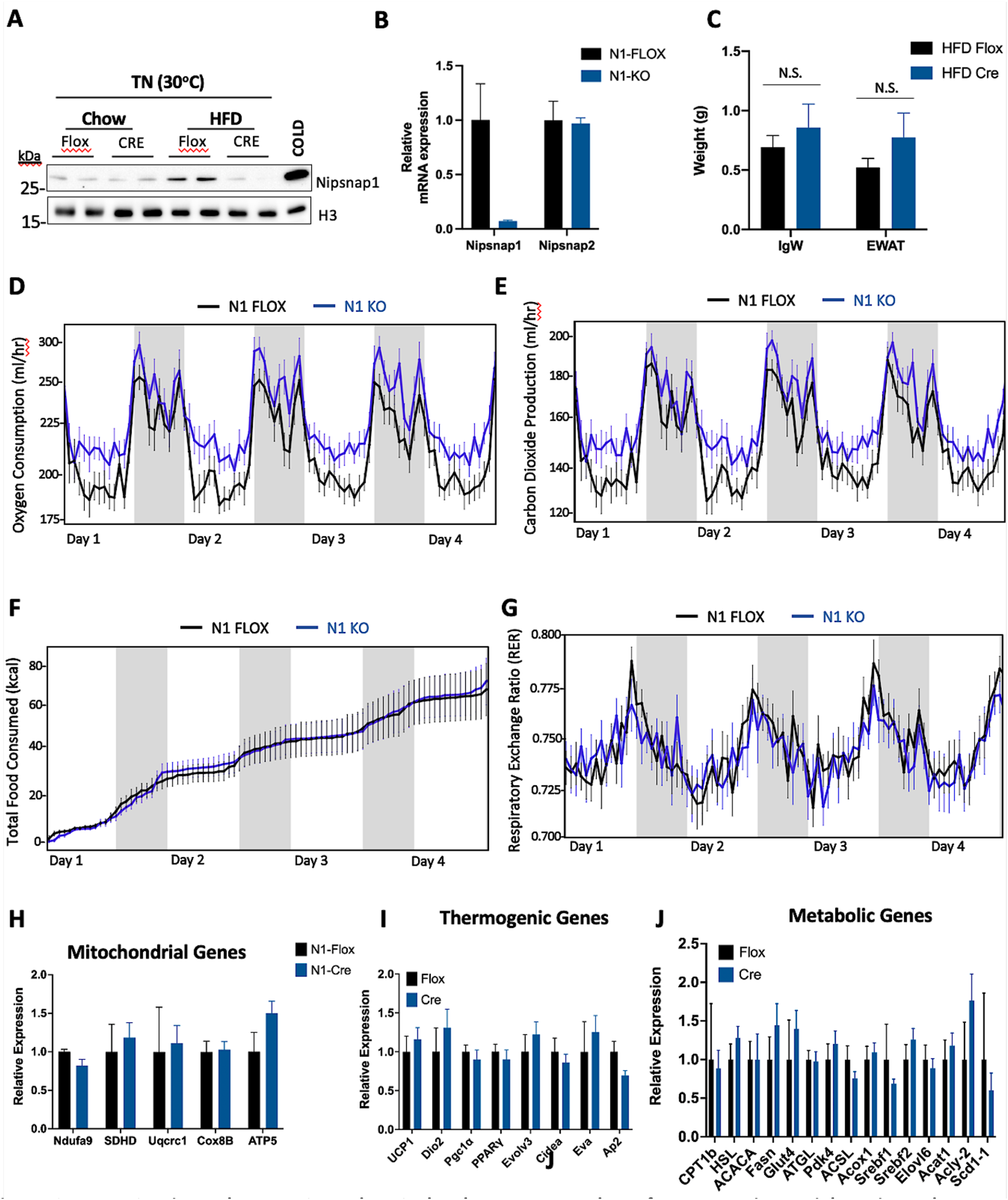
N1-KO mice enhance VO_2_ and VCO_2_ levels to prevent them from excessive weight gain under HFD conditions. (Related to Figure 4) **(A)**BAT representative Western blot of Nipsnap1 levels from chow and high-fat diet (HFD) fed N1-Flox and N1-KO mice for 24 weeks under thermoneutral conditions. COLD denotes N1-Flox mice exposed at cold (6°C) for 14 days as control. All experiments to follow are from N1-KO mice compared to N1-Flox controls fed a HFD regimen for 24 weeks under cold exposed conditions (6°C) (n=5) **(B)** Relative Nipsnap1 and Nipsnap2 mRNA levels **(C)** Tissue weights. **(D-G)** Oxygen consumption **(D)**, carbon dioxide generation **(E)**, food consumption **(F)**, and locomotive movement **(G)**. Grey bar indicates dark cycle and white bar indicates light cycle. **(H-J)** Relative mRNA expression of mitochondrial ETC (H), thermogenic (I) and lipid metabolic genes (J) in BAT tissue from N1-Flox and N1-KO mice exposed to high fat diet and cold exposure for 24 weeks (n=5). Whole body metabolic assessments **(D-G)** were analyzed by two-way ANOVA. All mRNA profiling was conducted by qPCR. mRNA profiling are represented as mean ± SEM. *p < 0.05, **p < 0.01 ***p < 0.001 by Student’s t test.

